# Active zone remodeling by Bruchpilot couples synaptic architecture to Kv1/Shaker excitability control

**DOI:** 10.64898/2025.12.26.696556

**Authors:** Sheng Huang, Chengji Piao, Marc Escher, David Toppe, Christine B. Beuschel, Torsten Götz, Niraja Ramesh, Zhiying Zhao, Oriane Turrel, Alexander M. Walter, Fan Liu, Stephan J. Sigrist

## Abstract

Presynaptic active zones are known to undergo state-dependent remodeling across sleep, circadian, and experience-dependent conditions, yet how such structural changes influence synaptic computation and excitability has remained unclear. Here, we address this gap by examining the functional consequences of physiological upscaling of the active zone scaffold Bruchpilot (BRP), within the range previously observed during natural state-dependent plasticity. We show that moderate BRP elevation expands the number of functional release sites while surprisingly reducing vesicle release probability, thereby establishing a presynaptic operating mode with selectively enhanced transmission at intermediate firing frequencies. This remodeled mode depends on Kv1/Shaker potassium channels, which normally constrain the increased structural capacity generated by BRP; accordingly, perturbation of Shaker abolishes BRP-dependent reductions in release probability and unmasks an enlarged synaptic output capacity. To test the functional relevance of this coupling, we examined *sleepless* mutants, in which Kv1/Shaker channels are destabilized and presynaptic remodeling is compromised. We show that direct, physiological-level BRP upscaling selectively restores the Shaker/Hyperkinetic channel complex from near-undetectable levels toward normal abundance without inducing global proteomic changes, and correspondingly rescues excitability balance, oxidative stress resistance, lifespan, and mid-term memory. Together, these findings identify a mechanistic coupling between active zone architecture and intrinsic excitability control and demonstrate how presynaptic structural plasticity shapes frequency-dependent transmission and functional robustness under stress.

## Introduction

Neural circuits must maintain functional stability despite ongoing challenges arising from metabolic demand, fluctuating network activity, sleep–wake transitions, and aging. To achieve such resilience, neurons employ complementary forms of homeostatic plasticity at intrinsic and synaptic levels. Changes in intrinsic excitability, including modulation of voltage-gated potassium channels such as Kv1/Shaker, critically regulate spike waveform, firing probability, and neurotransmitter release, thereby shaping network participation and protecting circuits against runaway excitation (*1–3*). Hyperexcitability is increasingly recognized as an early driver of pathological processes, including cognitive decline and neurodegenerative progression (*4*). In Drosophila, Kv1/Shaker channels and their auxiliary β-subunit Hyperkinetic are central regulators of neuronal excitability and sleep homeostasis (*5–7*). Loss of the GPI-anchored protein Sleepless/Quiver (SSS) destabilizes Shaker channels, producing severe hyperexcitability, reduced sleep, and shortened lifespan due to a physical loss of the Kv1/Shaker complex (*8, 9*).

Concurrently, recent work reveals that the presynaptic active zone (AZ) scaffold Bruchpilot (BRP), a core ELKS-family protein required for Ca²⁺ channel clustering and synaptic vesicle release (*10, 11*), is far more plastic and state-dependent than previously assumed. BRP abundance increases with sleep pressure and after sleep deprivation, forming a brain-wide presynaptic remodeling program termed PreScale that encodes accumulated sleep need (*12, 13*). Sleep deprivation also induces brain-wide BRP increases and active zone structural changes of cholinergic synapses (*14, 15*). Circadian systems regulate AZ morphology as well: CRY-dependent structural remodeling of visual system tetrads occurs at the morning activity peak (*16*), while circadian programming of the ellipsoid body influences sleep homeostat function and presynaptic output (*17, 18*). Beyond sleep and circadian control, BRP patterning is dynamically modulated by neuromodulators such as octopamine (*19*), and dopaminergic memory circuits engage AZ reorganization during active forgetting and individuality in sensory preference (*20, 21*). More broadly, AZ protein abundance, including BRP and Ca²⁺ channels, varies across inputs and contributes to functional synaptic diversity (*22*).

Despite this growing body of evidence, the functional significance of large-scale, state-dependent adjustments in BRP abundance and AZ architecture remains poorly understood. It is unknown whether these widespread presynaptic changes simply correlate with altered behavioral or physiological states or whether they actively contribute to synaptic computation, circuit stability, or resilience. In particular, the field lacks mechanistic insight into how structural remodeling at release sites interacts with intrinsic excitability mechanisms, such as Kv1-channel–mediated repolarization, and how whether these interactions might shape decoding of action potential patterns (via e.g. changes of short-term plasticity), or form in response to metabolic or behavioral demands. Thus, a central question emerges: what do brain-wide changes in active zone architecture actually *do* for synapses, circuits, and the organism?

Here, we address this question by dissecting the mechanistic, molecular, and behavioral consequences of moderate BRP upscaling, within the range naturally induced by sleep deprivation or early aging. We show that BRP elevation is associated with an apparent increase in functional release-site number, while unexpectedly, reducing vesicle release probability, yielding a presynaptic state optimized for frequency-selective transmission. This remodeled state critically depends on Kv1/Shaker channels. To test the functional relevance of this coupling under conditions of impaired excitability control, we examined *sleepless* (*sss*) mutants, which fail to engage presynaptic scaling. In this context, BRP upscaling selectively stabilizes the Shaker/Hyperkinetic channel complex, restoring its abundance and localization without inducing global proteomic restructuring. Supplying a physiological third brp copy bypasses the failure to engage presynaptic scaling in *sss* mutants and rescues synaptic hyperexcitability, oxidative stress sensitivity, lifespan, and mid-term memory, whereas identical BRP elevation does not rescue Shaker mutants.

Together, these findings identify a coupling between AZ remodeling and intrinsic excitability control, showing that BRP-dependent structural expansion interacts with Shaker-mediated repolarization to shape how synapses process activity. This mechanistic module provides a conceptual framework for how state-dependent AZ scaling can tune circuit function and contribute to neuronal and organismal resilience across sleep, circadian, and experience-dependent contexts.

## Results

Previous work has shown that ELKS-family scaffold Bruchpilot (BRP) abundance increases modestly under physiological conditions such as sleep deprivation and early aging, corresponding to an effective elevation from the endogenous two-copy state toward three genomic copies, whereas higher levels exceed this physiological range (*13, 15*). To determine how such state-dependent changes in active zone (AZ) architecture influence presynaptic function, we first took advantage of the Drosophila larval neuromuscular junction (NMJ), a glutamatergic synapse that permits high-resolution electrophysiological analysis of presynaptic release properties.

We thus titrated the copy number of *brp* from one to four genomic copies to systematically relate AZ scaling to presynaptic function. Increasing *brp* gene dosage produced a robust, monotonic rise in spontaneous miniature excitatory junctional current (mEJC) frequency (Fig. 1a). Because mEJC amplitudes and amplitude distributions remained unchanged across all genotypes (Fig. 1b,c), this suggests postsynaptic sensitivity was stable, indicating that the increase in spontaneous release reflects presynaptic changes. Under these conditions, the elevated mEJC frequency is consistent with an increase in the number of functional release sites, as inferred from spontaneous release measurements rather than direct structural quantification. Consistent with this interpretation, prior work has shown that the per AZ numbers of Unc13A clusters, which correlates with functional release-site number at the NMJ (*23*), increases with elevated BRP levels (*12*).

**Fig. 1.**
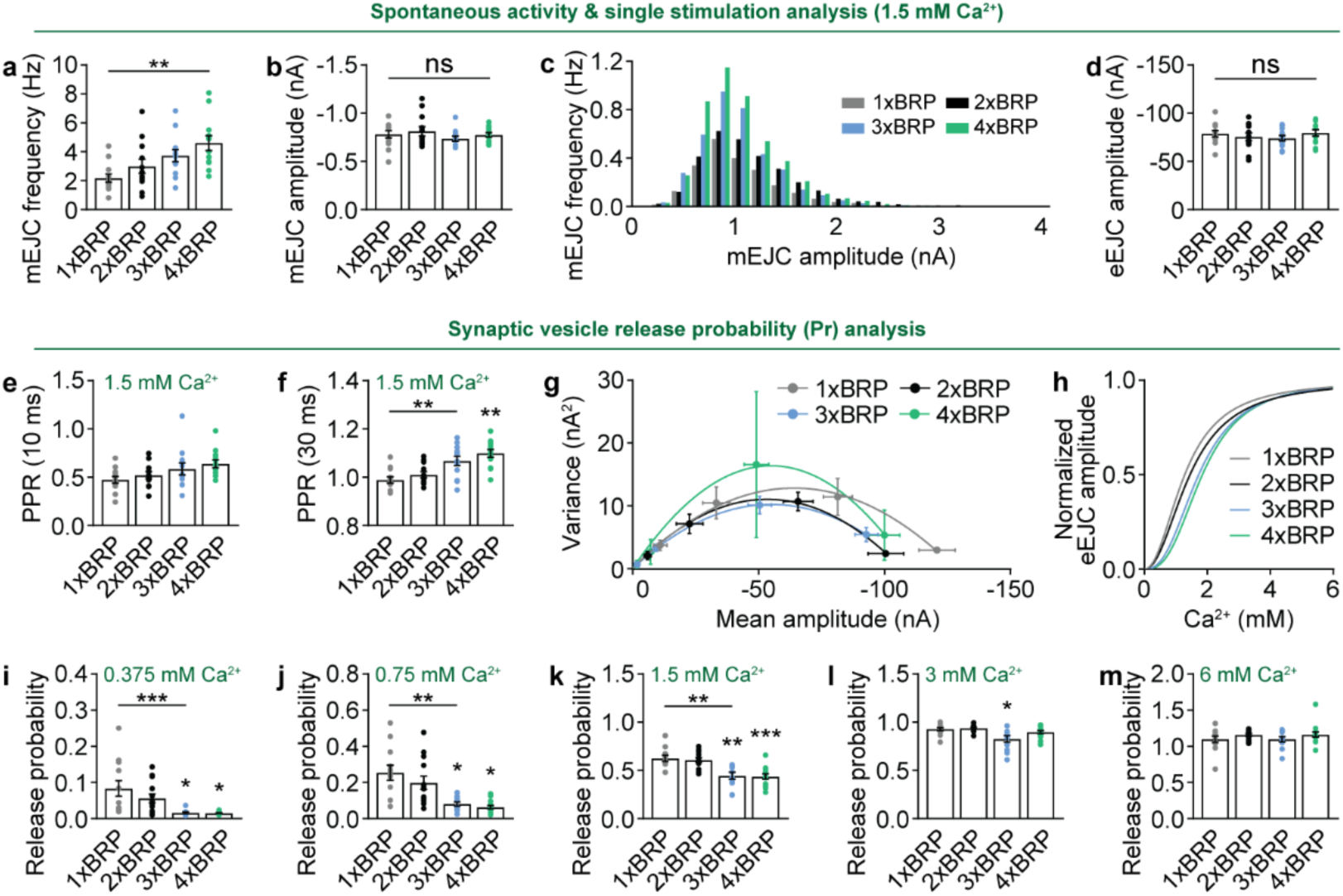
BRP-driven active zone remodeling dampens synaptic vesicle release probability. **a-c**, Spontaneous miniature excitatory junctional current (mEJC) analysis of the mean frequency, amplitude and distribution for 1x-4xBRP animals. n = 12. **d**, Evoked excitatory junctional current (eEJC) amplitude for 1x-4xBRP animals. n = 12. **e-f**, Pair-pulse ratio (PPR) analysis with inter-stimulation interval (ISI) of 10 ms or 30 ms for 1x-4xBRP animals. n = 12. **g-h**, Mean variance analysis for 1x-4xBRP animals. n = n = 11-15. **i-m**, Vesicle release probability at different Ca^2+^ concentration calculated from mean variance analysis for 1x-4xBRP animals. n = 11-15. \**p* < 0.05; \*\**p* < 0.01; \*\*\**p* < 0.001; ns, not significant. Error bars: mean ± SEM.

Despite the clear increase in release-site number implied by spontaneous transmission, single action potential–evoked eEJCs remained essentially unchanged between 1× and 4×BRP animals (Fig. 1d). Instead, our data reveal mechanistic differences that reflect distinct underlying modes by which neurotransmitter release is organized at the presynaptic terminal as BRP levels increase. Paired-pulse measurements showed a graded increase in facilitation with higher BRP copy number (Fig. 1e,f). This behavior strongly suggested a reduction in vesicle release probability (Pr) as BRP levels increased, an interpretation we tested directly using mean–variance analysis across a range of extracellular Ca²⁺ concentrations (Fig. 1g–m). Fitting parabolic variance–mean curves revealed a systematic reduction of Pr with each additional BRP copy, even at physiological Ca²⁺ (1.5 mM), with the effect only being overcome at supraphysiological concentrations (3–6 mM Ca²⁺). These findings suggest that BRP expansion increases the number of available release sites while simultaneously reducing the probability of release per site, thereby shifting synapses into a state with increased structural capacity and lower Pr.

### Shaker K⁺ channels gate BRP-dependent remodeling and constrain synaptic output

So far, our results suggest BRP upscaling as a presynaptic remodeling mechanism that regulates release statistics through coordinated changes in release-site abundance and vesicle release probability. By expanding structural release capacity while simultaneously lowering Pr, BRP upscaling might generates a presynaptic state in which synaptic output becomes particularly sensitive to factors that shape action potential waveform and Ca²⁺ influx. This organizational shift suggests that intrinsic excitability mechanisms may act downstream of, or in concert with, active zone remodeling to constrain the functional consequences of increased release capacity.

Shaker K⁺ channels gate BRP-dependent remodeling and constrain synaptic output Kv1-type Shaker channels are well positioned to fulfill such a role, as they accelerate presynaptic repolarization, limit Ca²⁺ entry, and thereby tune neurotransmitter release on a spike-by-spike basis. We therefore investigated whether Shaker activity interacts with BRP-dependent active zone remodeling to gate the functional expression of increased release-site number.

Acute pharmacological inhibition of Shaker with 4-aminopyridine (4-AP, 100 µM) caused a pronounced increase in evoked excitatory junctional current (eEJC) amplitudes across all BRP genotypes (Fig. 2a,b). Under these conditions, eEJC amplitudes scaled directly with BRP copy number, unmasking a latent synaptic capacity that is normally constrained by Shaker-mediated repolarization. This BRP-dependent facilitation under 4-AP was further reflected in prolonged decay constants and increased charge transfer during evoked responses (Fig. 2c,d).

**Fig. 2.**
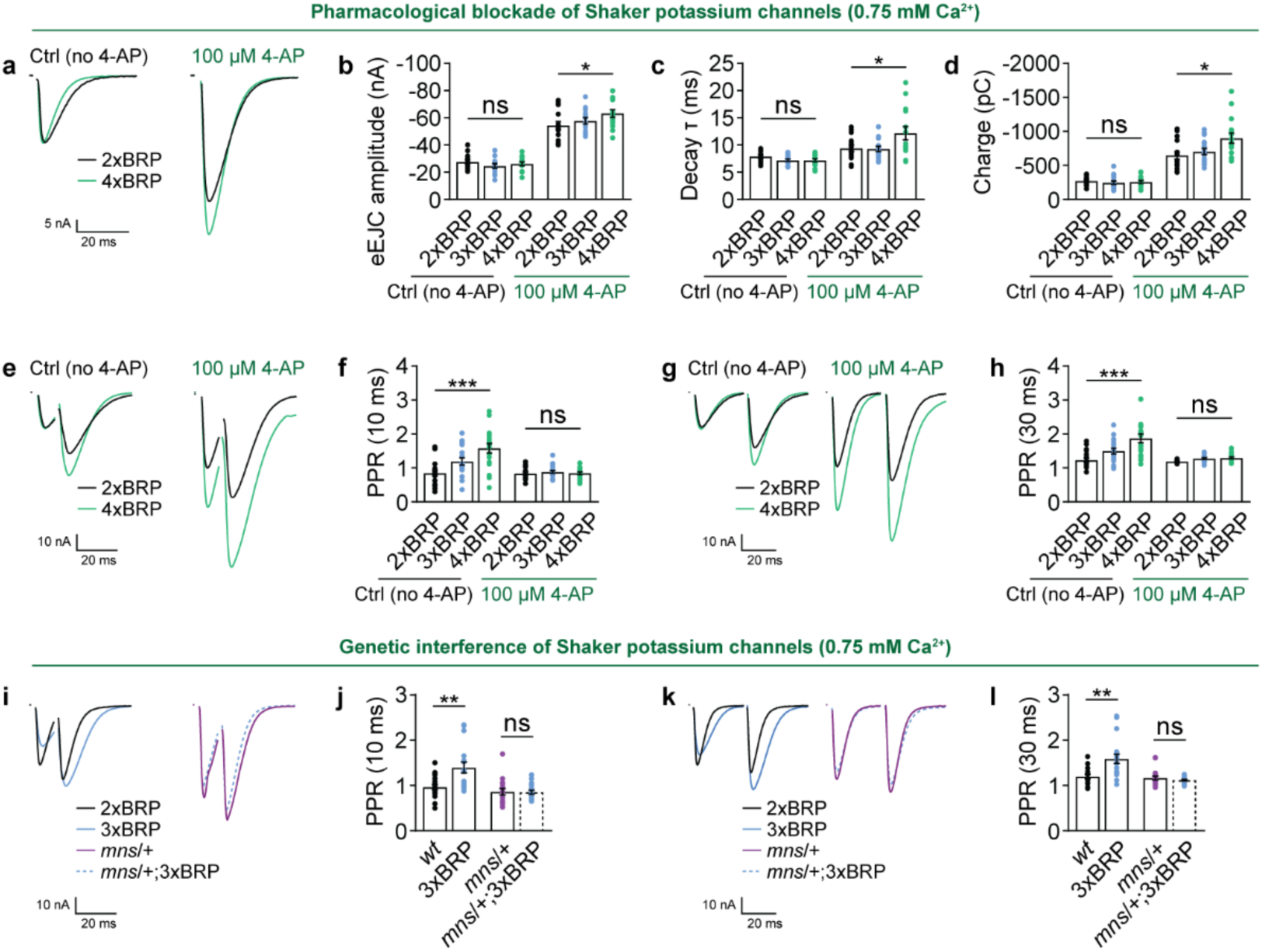
Shaker constrains BRP-driven active zone remodeling in promoting synaptic transmission. **a-d**, Representative eEJC traces, amplitudes, decay constants and charges for 2x-4xBRP animals with or without 100 μM 4-aminopyridine (4-AP) treatment. n = 13-14. **e-h**, Representative PPR traces of 10 ms and 30 ms ISI for 2x-4xBRP animals with or without 100 μM 4-AP treatment. n = 17-18. **i-l**, Representative PPR traces of 10 ms and 30 ms ISI for 2x-3xBRP animals in *mns*/+ or *wt* background. n = 18. \**p* < 0.05; \*\**p* < 0.01; \*\*\**p* < 0.001; ns, not significant. Error bars: mean ± SEM.

Notably, 4-AP also eliminated BRP-dependent differences in paired-pulse ratios (Fig. 2e–h). Whereas elevated BRP normally reduces Pr, blocking Shaker rendered synapses from different BRP genotypes indistinguishable in their short-term plasticity. These results indicate that Shaker activity is required for the BRP-dependent reduction of release probability and that the suppressive action of Shaker masks the expanded release capacity produced by AZ remodeling.

To exclude off-target pharmacological effects, we next introduced a heterozygous Shaker loss-of-function mutation (*mns*/+) into a 3×BRP background. Genetic reduction of Shaker phenocopied the effects of 4-AP: BRP-dependent differences in PPR were eliminated (Fig. 2i–l). Together, these pharmacological and genetic manipulations indicate that Shaker channels function as physiological constraints on BRP-driven remodeling, potentially by limiting Ca²⁺ influx and preserving synaptic stability in the face of increased release-site number.

Thus, BRP and Shaker jointly define a two-component system for presynaptic tuning: BRP expands the structural capacity of the AZ, whereas Shaker acts as a brake that prevents excessive synaptic output. This mechanistic coupling provides the basis for frequency-dependent modulation.

### BRP and Shaker generate a frequency-selective transmission filter at NMJ synapses

We next asked how this combined BRP–Shaker module modulates transmission across different firing regimes. Under high-frequency stimulation (100 Hz), cumulative quantal content, readily releasable pool (RRP) size, and vesicle replenishment rates were indistinguishable between BRP genotypes (Fig. 3a–c). These data suggest that under extreme activity, processes downstream of Ca²⁺ influx, such as vesicle mobilization and RRP exhaustion, become rate-limiting, diminishing the influence of AZ remodeling.

**Fig. 3.**
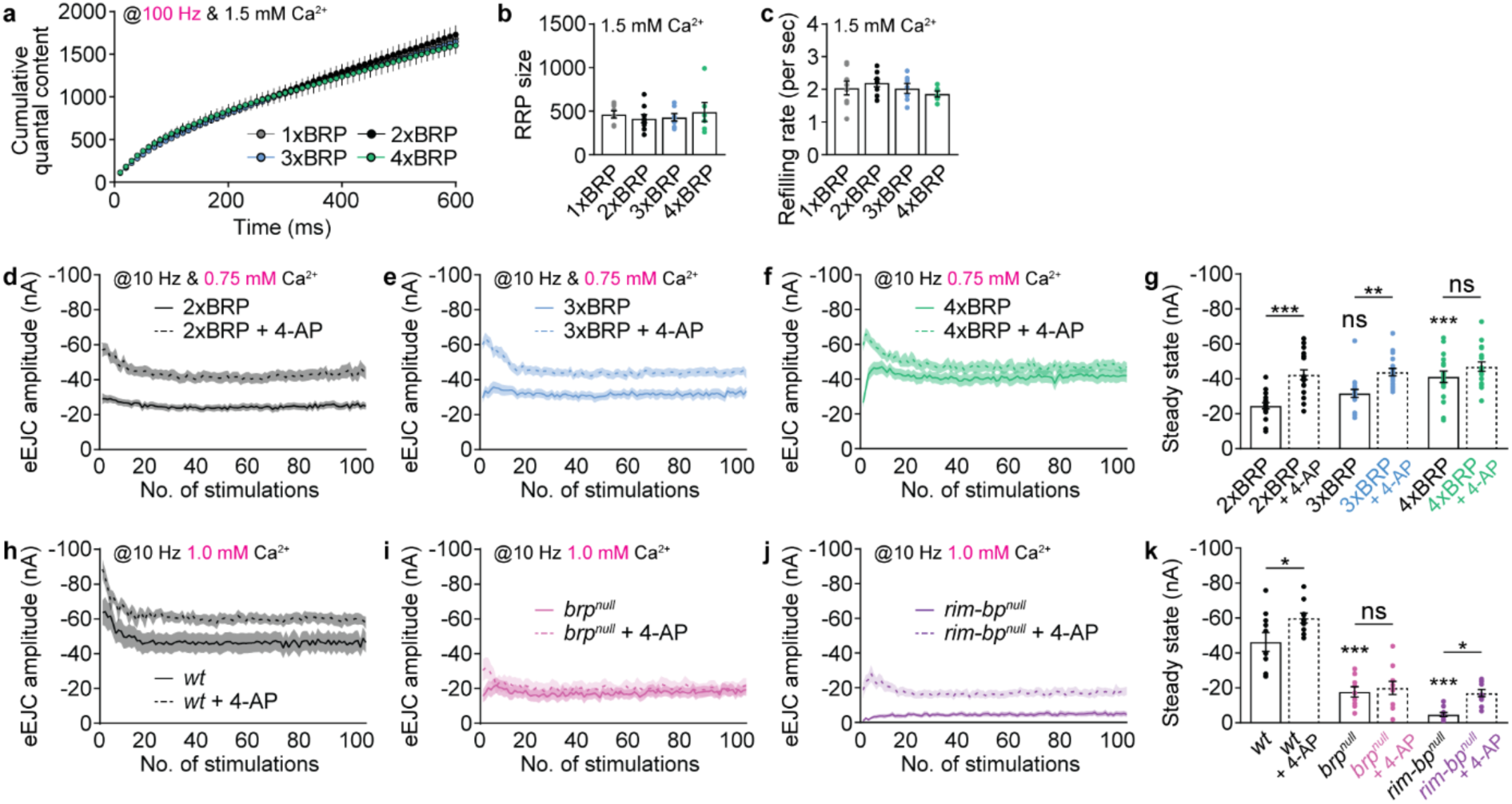
BRP-driven active zone remodeling promotes synaptic transmission in a frequency-dependent manner. **a-c**, Cumulative quantal content, readily releasable pool (RRP) and refilling rate for 1x-4xBRP animals. n = 6-9. **d-g**, Mean eEJC and plateau amplitudes for 2x-4xBRP animals with or without 100 μM 4-AP treatment stimulated at 10 Hz. n = 17-18. **h-k**, Mean eEJC and plateau amplitudes for *wt*, *brp* null and *rim-bp* null animals with or without 100 μM 4-AP treatment stimulated at 10 Hz. n = 9-10. \**p* < 0.05; \*\**p* < 0.01; \*\*\**p* < 0.001; ns, not significant. Error bars: mean ± SEM.

In contrast, at intermediate frequencies (10 Hz), BRP copy number strongly influenced steady-state synaptic output. Higher BRP levels produced a robust enhancement of 10 Hz transmission (Fig. 3d–g), revealing a frequency-selective facilitation that did not manifest at low (0.2 Hz) or high (100 Hz) frequencies. This finding indicates that BRP remodeling tunes synapses to operate optimally within a distinct physiological range.

The Shaker dependence of this effect was again evident: 4-AP application increased 10 Hz transmission in 2×BRP synapses but failed to further enhance output in 4×BRP animals (Fig. 3d–g), suggesting that BRP upscaling and Shaker blockade act on the same regulatory mechanism and converge on a shared ceiling of Ca²⁺ influx. To test whether other AZ components similarly gate Shaker effects, we compared *brp* null and *rim-bp* null mutants. Strikingly, rim-bp mutants showed increased 10 Hz transmission upon 4-AP treatment, whereas brp mutants did not (Fig. 3h–k). This specificity argues that BRP, not RIM-BP, is the critical molecular platform for Shaker-dependent excitability control at the AZ.

Collectively, these results demonstrate that BRP and Shaker jointly implement a presynaptic excitability filter that reduces release at low frequencies, enhances throughput at intermediate frequencies, and becomes saturated at high frequencies. Such frequency-selective tuning provides a plausible mechanism by which neurons could adapt synaptic output to physiological demand while minimizing metabolic stress.

### BRP upscaling restores excitability control in *sleepless* mutants

Having established a functional interaction between BRP and the Shaker potassium channel, we next asked whether BRP directly influences Shaker protein abundance or subcellular localization. To address this, we used pan-neuronal expression of a Shaker-RFP fusion protein and compared Shaker levels in brains carrying either two or three genomic copies of brp. Strikingly, Shaker protein levels were significantly elevated in 3×BRP compared with 2×BRP brains (Fig. 4a,b). Because the Shaker cDNA was expressed from the same chromosomal integration site in all genotypes, differences in Shaker protein levels cannot be attributed to transcriptional regulation, indicating that BRP promotes Shaker stability through post-transcriptional mechanisms. This interpretation is consistent with earlier observations in *sleepless* (*sss*) mutants, in which Shaker protein levels are reduced despite unaltered transcript abundance, pointing to regulation at the level of protein stability or turnover.

**Fig. 4.**
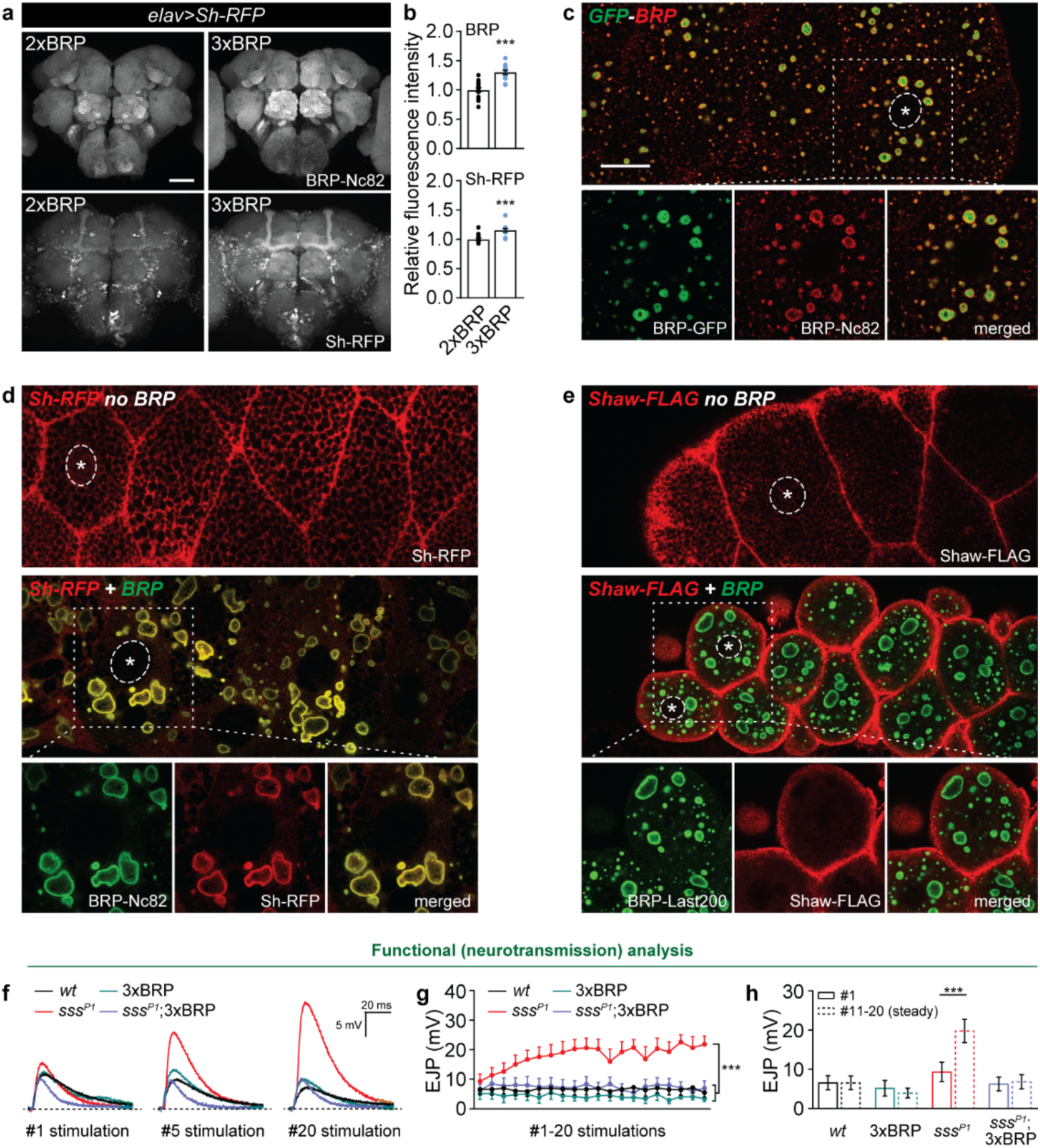
BRP promotes Shaker level and potentially directs its subcellular localization. **a-b**, Representative confocal images and whole-mount brain staining analysis of BRP and *elav-Gal4*-driven Shaker-RFP expression levels in 2xBRP and 3xBRP brains. n = 13-15. Scale bar: 50 μm. **c**, Representative confocal images of salivary glands in *OK6-Gal4*-driven expression of GFP-tagged BRP. Anti GFP and BRP (Nc82) immunostaining are shown. **d**, Representative confocal images of salivary glands in *OK6-Gal4*-driven expression of RFP-tagged Shaker with or without the co-expression of BRP. Anti RFP and BRP (Nc82) immunostaining are shown. **e**, Representative confocal images of salivary glands in *OK6-Gal4*-driven expression of FLAG-tagged Shaw with or without the co-expression of BRP. Anti FLAG and BRP (Last200) immunostaining are shown. Scale bar: 25 μm. **f-h**, Current clamp showing the evoked excitatory junctional potential (eEJP) augmentation phenotype of *sss^P1^* and suppression by one more gene copy of BRP. Stimulation was performed at 0.2 Hz frequency. n = 8. \**p* < 0.05; \*\**p* < 0.01; \*\*\**p* < 0.001; ns, not significant. Error bars: mean ± SEM.

To test whether BRP might directly influence Shaker trafficking or spatial targeting, we turned to a heterologous, non-neuronal system. Ectopic expression of BRP in larval salivary glands resulted in the formation of large intracellular, condensate-like structures (Fig. 4c), consistent with the known propensity of BRP to self-assemble into scaffolded compartments. When Shaker-RFP was co-expressed in this system, Shaker was robustly recruited into these BRP-positive compartments. In contrast, the closely related voltage-gated potassium channel Shaw showed no detectable enrichment in BRP condensates (Fig. 4d,e). This selective recruitment indicates molecular specificity and supports the idea that BRP scaffolds can directly organize Shaker-containing channel complexes, rather than acting through a generic effect on membrane proteins.

We next asked whether the BRP-dependent effects on Shaker stability described above have functional consequences for neuronal excitability in vivo. Loss of the GPI-anchored protein Sleepless/Quiver (SSS) destabilizes the Kv1/Shaker channel complex, resulting in pronounced neuronal hyperexcitability, reduced sleep, and shortened lifespan. At the molecular level, this phenotype reflects a physical loss of Shaker channels, which is readily detectable by immunostaining and Western blot analysis (*8, 9*).

Importantly, unlike other short-sleep *Drosophila* mutants, *sleepless* mutant brains fail to engage a compensatory upscaling of BRP (Fig. 1 in (*12*)). This selective absence of BRP upregulation suggested that *sss* mutants might lack a homeostatic buffering mechanism normally recruited under conditions of elevated excitability. We therefore used *sleepless* mutants as a sensitized genetic background to test whether experimentally elevating BRP levels is sufficient to restore excitability control and synaptic output stability in the context of impaired Shaker channel function.

Consistent with previous reports, *sss* larvae exhibited a progressive increase in excitatory junctional potential amplitude during low-frequency (0.2 Hz) stimulation, a hallmark of defective excitability control (Fig. 4f–h). Remarkably, introduction of a third *brp* copy fully suppressed this phenotype, restoring synaptic output dynamics to wild-type levels. Together, these findings indicate that BRP upscaling can compensate for the excitability defects of *sss* mutants by stabilizing Shaker abundance and function, thereby re-establishing effective control over synaptic excitability. We wondered whether this functional rescue might be due to a stabilization of the Shaker complex in this constellation.

### BRP upscaling restores the Shaker/Hyperkinetic complex in *sss* mutant brains

To investigate this molecular rescue in more detail, we performed quantitative proteomics on dissected adult brains from control and sss backgrounds with either 2×BRP or 3×BRP. Principal component analysis revealed distinct separation of wild-type and sss samples but showed that 3×BRP did not broadly reconfigure the sss proteome (Fig. 5a). Heatmap analysis similarly indicated that BRP upscaling does not globally normalize the proteome (Fig. 5b), implying a targeted rather than “broadband” rescue.

**Fig. 5.**
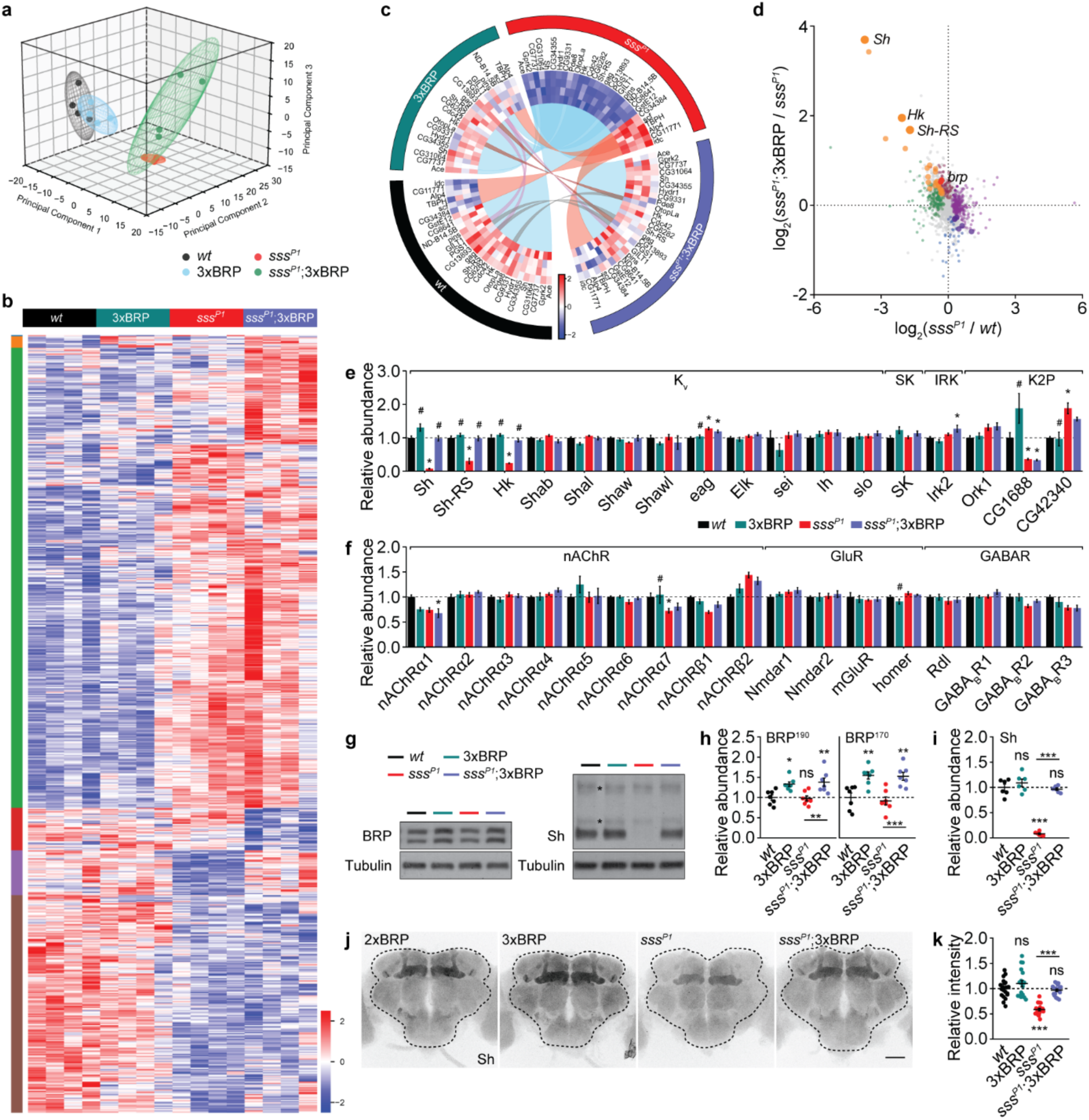
A genetic active zone upscaling restores voltage-gated Shaker K^+^ channel and its β subunit Hyperkinetic. **a**, Principal component analysis (PCA) for 2xBRP and 3xBRP in control or *sss^P1^* background. **b**, Heatmap of significantly regulated proteins for 2xBRP and 3xBRP in control or *sss^P1^* background. **c**, Circular heatmap showing proteins that exhibit significant interaction effects between the two factors (SSS and BRP). **d**, Scatter plot of LFQ MS data plotting the difference in protein levels induced by loss of SSS or by increasing BRP copies from 2 to 3. X-axis shows the log2 fold-change values representing the effects of *sss^P1^*, Y-axis displays the log2 of the p-values representing the effects of 3xBRP. **e**, Normalized protein abundance for detected K^+^ channels in control or *sss^P1^* mutants with either 2xBRP or 3xBRP. * indicates significant difference to *wt*. # indicates significant difference to *sss^P1^*. **f**, Normalized protein abundance for detected neurotransmitter receptors in control or *sss^P1^* mutants with either 2xBRP or 3xBRP. * indicates significant difference to *wt*. # indicates significant difference to *sss^P1^*. **g-i**, Representative Western blots and statistics of BRP and Shaker levels for 2xBRP and 3xBRP in control or *sss^P1^*background. **j-k**, Confocal images and whole-mount brain staining analysis of BRP and Shaker levels for 2xBRP and 3xBRP in control or *sss^P1^*mutant background. Scale bar: 50 μm. */# *p* < 0.05; \*\**p* < 0.01; \*\*\**p* < 0.001; ns, not significant. Error bars: mean ± SEM.

Remarkably, the Shaker/Hyperkinetic channel complex was selectively restored by BRP elevation in our brain proteomic data. Whereas Shaker and Hk levels were almost undetectable in *sss^P1^,2×BRP brains*, their abundance in *sss^P1^;3×BRP* animals approached wild-type levels (Fig. 5c–f). Western blotting confirmed this restoration (Fig. 5g–i), and whole-brain immunostaining revealed normalization of Shaker localization, prominently in mushroom body axons (Fig. 5j,k). Additional analyses using the milder *sss^P2^* allele confirmed consistent rescue (Extended Data Fig. 1a–d), while higher BRP (4×) further increased Shaker abundance (Extended Data Fig. 1e,f), suggesting a dosage-sensitive relationship.

Thus, BRP elevation restores specific molecular machinery of intrinsic excitability in *sss* mutants without broadly altering protein expression, identifying the Shaker/Hk complex as a specific and sensitive substrate of PreScale remodeling.

### BRP-mediated restoration of the Shaker complex rescues resilience, memory, and lifespan in *sleepless* mutants

We finally assessed whether this molecular and synaptic rescue translates into organismal resilience. As expected, increasing *brp* gene copy number had little effect on H₂O₂ sensitivity or lifespan in wild-type animals (Fig. 6a,b). In contrast, *sss^P1^,2×BRP* flies exhibited profound oxidative sensitivity and greatly reduced lifespan, both of which were robustly restored by adding a third *brp* copy (Fig. 6c,d). In the *Shaker^mns^* mutants, however, the same BRP elevation failed to rescue oxidative stress sensitivity or lifespan and even exacerbated mortality (Fig. 6e,f), demonstrating that BRP exerts its protective effect through Shaker.

**Fig. 6.**
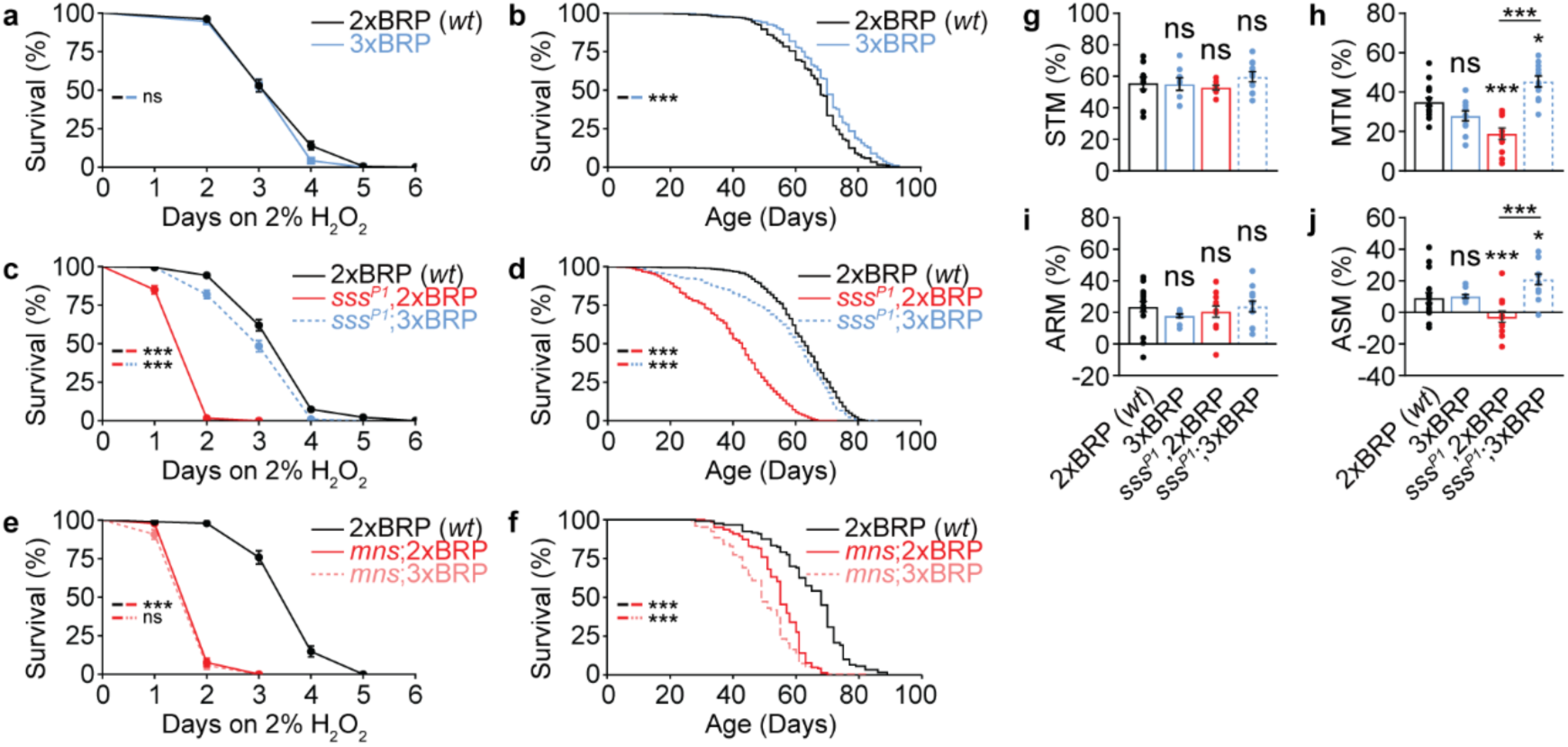
Elevating *brp* copy number restores survival and olfactory aversive mid-term memory in *sss* mutants. **a**, Survival curves for 2xBRP *wt* and 3xBRP flies under 2% H_2_O_2_ treatment. n = 157-165. **b**, Survival curves showing the lifespan of 2xBRP *wt* and 3xBRP flies. n = 475-483. **c**, Survival curves for *wt* control and *sss^P1^* in either 2xBRP or 3xBRP background under 2% H_2_O_2_ treatment. n = 176-182. **d**, Survival curves showing the lifespan of *wt* control and *sss^P1^*in either 2xBRP or 3xBRP background. n = 826-832. **e**, Survival curves for *wt* control and *Shaker mns* in either 2xBRP or 3xBRP background under 2% H_2_O_2_ treatment. n = 86-95. **f**, Survival curves showing the lifespan of *wt* control and *Shaker mns* in either 2xBRP or 3xBRP background. n = 120-146. **g**, Short-term memory (STM) for 2xBRP and 3xBRP in *wt* control or *sss^P1^* background. n = 8-10. **h**, Middle-term memory (MTM) for 2xBRP and 3xBRP in *wt* control or *sss^P1^* background. n = 11-16. **i**, Anesthesia-resistant memory (ARM) for 2xBRP and 3xBRP in *wt* control or *sss^P1^* background. n = 11-17. **j**, Anesthesia-sensitive memory (ASM) for 2xBRP and 3xBRP in *wt* control or *sss^P1^* background. n = 11-17. \**p* < 0.05; \*\**p* < 0.01; \*\*\**p* < 0.001; ns, not significant. Error bars: mean ± SEM.

We next examined cognitive performance. Short-term memory remained unaffected across genotypes (Fig. 6g), but mid-term memory (MTM) was severely impaired in *sss^P1^,2×BRP* animals and fully restored in *sss^P1^;3×BRP flies* (Fig. 6h). MTM consists of anesthesia-resistant memory (ARM) and anesthesia-sensitive memory (ASM). ARM remained unchanged (Fig. 6i), whereas ASM (was abolished in *sss*) was significantly enhanced above control levels in *sss^P1^;3×BRP* animals (Fig. 6j).

We also examined the consequences of excessive BRP upscaling. In wild-type flies, 4×BRP significantly impaired memory performance and reduced survival (Extended Data Fig. 2a–c). In *sss* mutants, 4×BRP further deteriorated their already compromised phenotypes. These findings underscore the importance of keeping the BRP regulation in a physiological range and support the view that PreScale reflects a tightly tuned homeostatic response.

Sleep analyses showed partial restoration of nighttime sleep in sssP1;3×BRP animals (Extended Data Fig. 3a). However, sleep improvement cannot fully explain the resilience rescue: in *Shaker^mns^* mutants, increasing BRP also increased sleep (Extended Data Fig. 3b), yet did not rescue stress sensitivity or lifespan (Fig. 6e,f). Thus, restored excitability control— not sleep—is the primary driver of resilience.

Together, these results demonstrate that bypassing the failure to induce elevated BRP levels in *sss* mutants, achieved through BRP upscaling, reinstates Shaker abundance, rescues synaptic excitability, and restores organismal resilience across oxidative, cognitive, and lifespan metrics.

## Discussion

Neuronal circuits must maintain stable function despite continuous challenges imposed by fluctuating metabolic demand, variable activity patterns, and environmental stress. Achieving such robustness requires coordinated homeostatic mechanisms operating at both intrinsic and synaptic levels (*1*). Disruption of this balance, particularly in the form of neuronal hyperexcitability, is increasingly recognized as a driver of oxidative stress, impaired cognition, and reduced lifespan (*4*). Among the intrinsic mechanisms controlling excitability, Kv1/Shaker-class potassium channels play a central role by shaping action potential waveform and constraining presynaptic Ca²⁺ influx (*24, 25*). Although intrinsic excitability regulation and synaptic plasticity have long been studied in parallel (*26*), how structural remodeling of presynaptic release sites interfaces with ion-channel–based excitability control has remained poorly understood.

Here, we identify a mechanistic coupling between presynaptic active zone (AZ) architecture and intrinsic excitability regulation. Moderate elevation of the AZ scaffold Bruchpilot (BRP), within a physiological range previously observed across multiple behavioral and aging-related contexts (see introduction), induces a coordinated remodeling of presynaptic function. We find that BRP elevation increases the apparent number of functional release sites while simultaneously reducing vesicle release probability, thereby redistributing neurotransmitter release across a larger structural pool. This reorganization might establish a presynaptic operating mode that favors transmission at intermediate firing frequencies, without enhancing output at low or saturating frequencies. Importantly, this remodeled state critically depends on Kv1/Shaker potassium channels, which act to constrain the functional consequences of increased release capacity.

Our data support a model in which BRP-dependent expansion of release-site number sensitizes synaptic output to mechanisms that shape action potential waveform and Ca²⁺ influx. Shaker channels are ideally positioned to fulfill this role, as they accelerate repolarization and limit presynaptic Ca²⁺ entry on a spike-by-spike basis (*3, 24*). Pharmacological or genetic reduction of Shaker function unmasks a latent BRP-dependent increase in synaptic output, eliminating differences in short-term plasticity across BRP levels and revealing a direct scaling of evoked responses with BRP copy number. Thus, BRP and Shaker together seemingly define a two-component presynaptic system: BRP expands structural capacity, while Shaker provides a functional brake that preserves synaptic stability. At the neuromuscular junction, this interaction gives rise to a frequency-selective mode of transmission, implemented through coordinated changes in release-site number, vesicle release probability, and Shaker-dependent repolarization. One likely scenario is that at low frequencies, reduced release probability limits output; at intermediate frequencies, BRP-dependent expansion enables enhanced throughput; and at high frequencies, downstream constraints such as vesicle depletion dominate. Such frequency dependence may be particularly well suited to circuits operating within defined physiological firing regimes, allowing synapses to enhance information transfer without incurring excessive Ca²⁺ load or metabolic cost. In this view, AZ remodeling does not act as a simple gain increase but rather reconfigures how synapses transform activity patterns into neurotransmitter release.

A central finding of this study is that BRP elevation influences not only release machinery but also the stability of intrinsic excitability components. Increasing BRP levels selectively enhances the abundance of the Shaker/Hyperkinetic (Hk) channel complex without inducing broad or compensatory changes in our quantitative proteomics measurements (*27, 28*) of Drosophila brain proteomes. Notably, neither other potassium channels nor neurotransmitter receptors showed comparable restoration or modulation, underscoring the selectivity of this effect. This specific interaction is evident both in neurons and in a heterologous system, where BRP forms condensate-like assemblies that selectively recruit Shaker but not related potassium channels. While our data demonstrate that BRP elevation is sufficient to restore Shaker/Hk abundance and function in sensitized backgrounds, they do not yet resolve whether this stabilization reflects direct molecular interaction, altered trafficking, or changes in local retention or turnover.

The functional relevance of this coupling becomes particularly apparent in the sensitized background of *sleepless* (*sss*) mutants. Loss of the GPI-anchored protein Sleepless destabilizes the Kv1/Shaker complex, resulting in severe hyperexcitability and systemic vulnerability (*8, 9*). Notably, sss mutants fail to engage a compensatory elevation of BRP (*12*), suggesting a deficit in an otherwise available buffering mechanism. Supplying a physiological third copy of *brp* bypasses this failure and restores excitability control at synapses, normalizes oxidative stress resistance and lifespan, and rescues mid-term memory. The failure of identical BRP elevation to rescue Shaker mutants argues strongly against indirect effects mediated by development, sleep, or general neuronal health and demonstrate that the protective effects of BRP elevation depend on functional Shaker channels.

These findings indicate that BRP elevation can function as a compensatory response to impaired excitability control, provided that the underlying channel machinery remains responsive. At the same time, our data underscore the importance of tight quantitative regulation: excessive BRP elevation is detrimental, impairing memory and survival even in otherwise wild-type animals (*13*). Thus, AZ remodeling seemingly has to operate within a rather narrow physiological window, consistent with homeostatic models in which both insufficient and excessive compensation are harmful (*29*).

More broadly, our results provide a mechanistic framework for understanding why presynaptic architecture is dynamically regulated across diverse physiological conditions, including aging, neuromodulatory state changes, circadian transitions, and sleep-related contexts (*14, 16, 17, 19, 21*). Rather than serving merely as a correlate of these states, BRP-dependent remodeling might be understood as a means to reconfigure synaptic computation itself, expanding structural capacity while relying on intrinsic excitability mechanisms to prevent destabilization. By stabilizing the Shaker/Hyperkinetic complex, BRP elevation further reinforces this balance, linking synaptic structure directly to electrical control.

Several independent lines of work place our findings into a broader physiological context. Metabolic and oxidative states have been shown to directly modulate intrinsic excitability through the Shaker/Hyperkinetic complex, with redox-dependent regulation of Hyperkinetic altering Kv1 channel kinetics and neuronal firing under elevated oxidative load (*7, 30*). As mentioned above, conditions associated with prolonged wakefulness are accompanied by elevated BRP abundance, while our data indicate that elevated BRP is associated with reduced presynaptic release probability. Notably, recent preprint work further supports the functional relevance of a selective reduction in release probability itself as a determinant of extended wake phases (bioRxiv: 2025.10.16.682770). Together, these observations converge on a model in which metabolic and excitability stress engage a presynaptic remodeling response characterized by BRP elevation, constrained release probability, and stabilization of Kv1/Shaker-mediated repolarization—closely matching the configuration we identify as supporting frequency-selective transmission and synaptic robustness.

Several important questions remain. The molecular basis by which BRP stabilizes the Shaker/Hk complex, whether through direct interaction, shared trafficking pathways, or nanoscale compartmentalization, remains to be elucidated. The neuromuscular junction provides a uniquely well-resolved system to dissect presynaptic mechanisms; extending these principles to central synapses will be essential to establish their generality across circuit architectures. It will also be important to determine whether other AZ components engage similar coupling to intrinsic excitability regulators, such as RIM-BP or Unc13 isoforms

In summary, this study identifies a structural–electrical coupling mechanism at presynaptic terminals in which elevation of the active zone scaffold BRP expands release capacity while Shaker-dependent repolarization constrains excitability. Through this interaction, synapses acquire frequency-dependent filtering properties and enhanced robustness under conditions of excitability stress. These findings provide a conceptual framework for how dynamic regulation of presynaptic architecture can contribute to stabilizing neural function without sacrificing computational flexibility.

## FIGURES AND FIGURE LEGENDS

**Extended Data Fig. 1.**
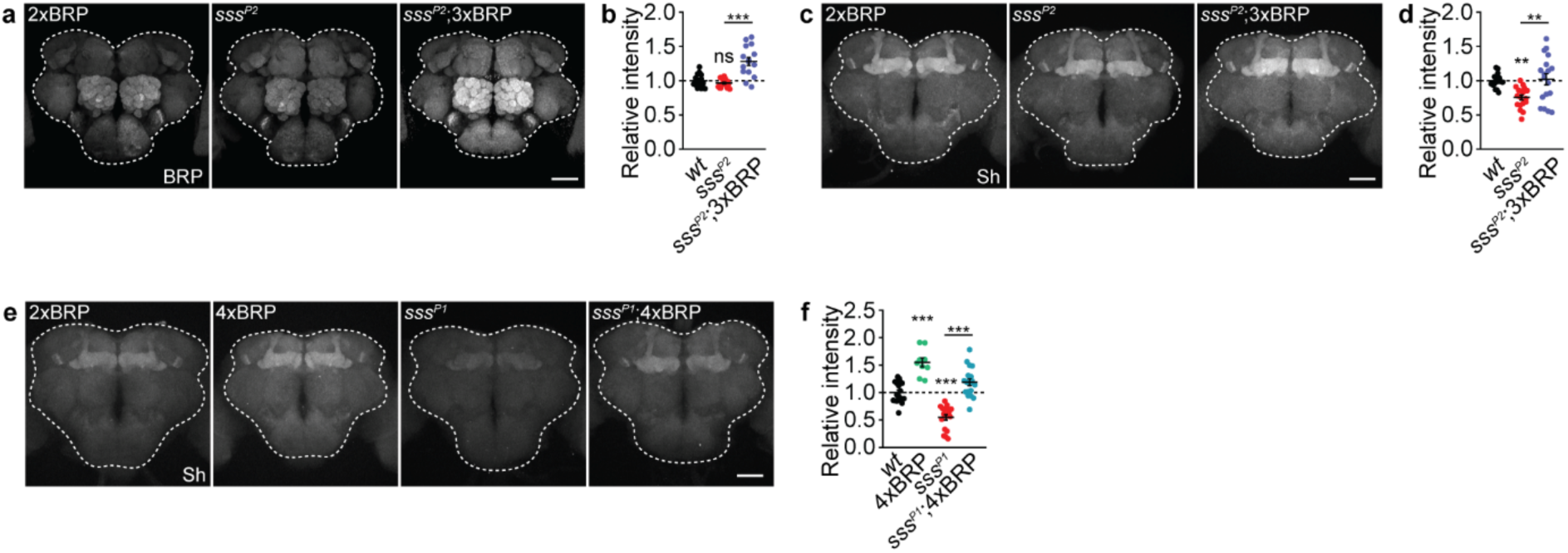
BRP-driven active zone remodeling restores Shaker level in both *sss^P1^* and *sss^P2^* mutants. **a-d**, Confocal images and whole-mount brain staining quantification of BRP and Shaker levels for 2xBRP and 3xBRP in control or *sss^P2^* mutant background. n = 17-18. Scale bar: 50 μm. **e-f**, Confocal images and whole-mount brain staining quantification of Shaker level for 2xBRP and 4xBRP in control or *sss^P1^* mutant background. n = 9-18. Scale bar: 50 μm. \*\**p* < 0.01; \*\*\**p* < 0.001; ns, not significant. Error bars: mean ± SEM.

**Extended Data Fig. 2.**
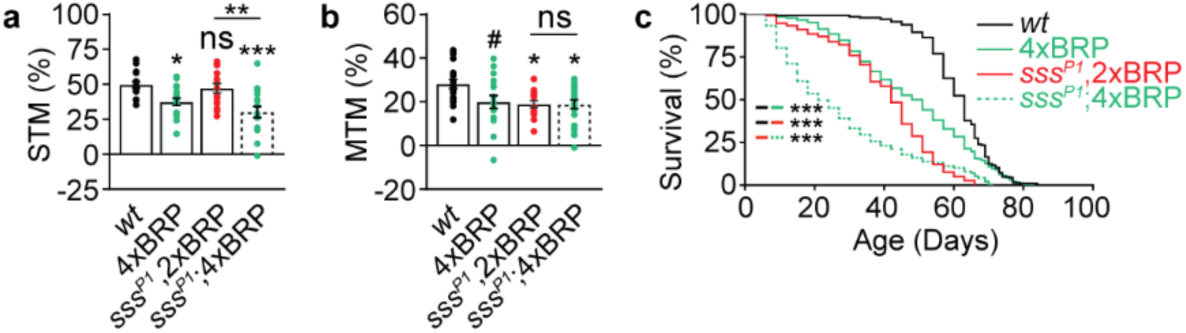
Excessive BRP-driven active zone remodeling is detrimental. **a-b**, Short-term memory (STM) and middle-term memory (MTM) for 2xBRP and 4xBRP in control or *sss^P1^* background. n = 15-21. **c**, Survival curves for lifespan of *wt* control and *sss^P1^* in either 2xBRP or 3xBRP background. n = 223-308. \**p* < 0.05; \*\**p* < 0.01; \*\*\**p* < 0.001; ns, not significant. Error bars: mean ± SEM.

**Extended Data Fig. 3.**
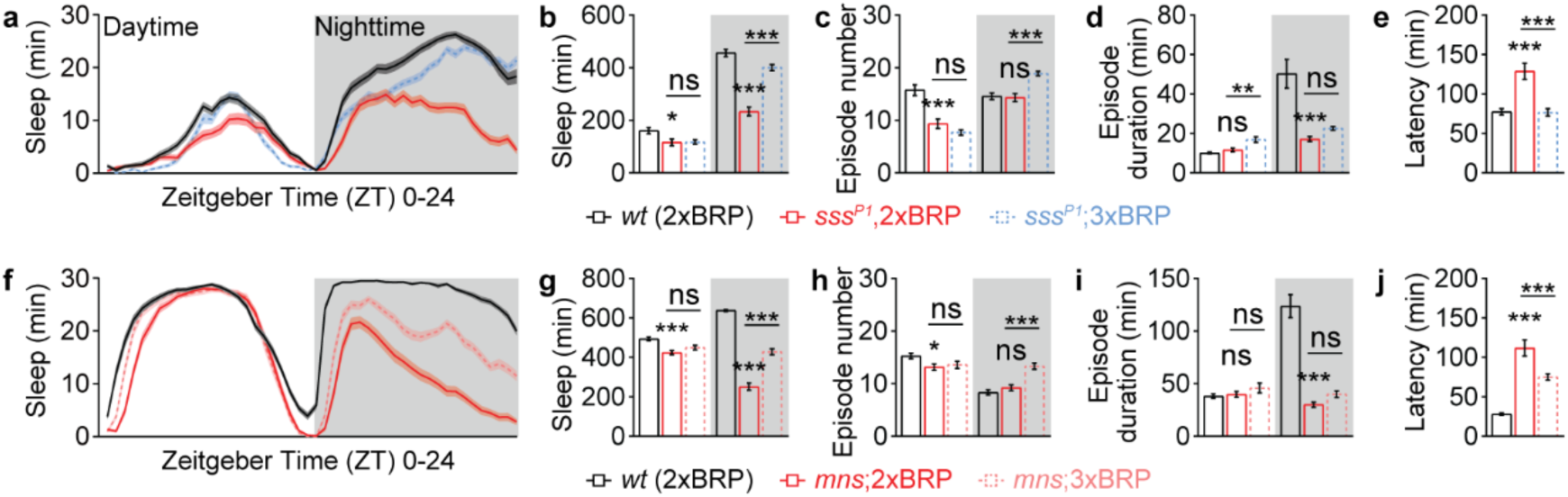
BRP-driven active zone remodeling promotes sleep in both *sss^P1^* and *Shaker mns* mutants. **a-e**, Sleep profile of 2xBRP *wt*, *sss^P1^*,2xBRP and *sss^P1^*;3xBRP averaged from measurements over 2-4 days, including sleep curves plotted in 30-min bins, daily sleep amount, sleep episode number, sleep episode duration and sleep latency at ZT12. Data from female flies are shown. n= 48-72. **a-e**, Sleep profile of 2xBRP *wt*, *mns*;2xBRP and *mns*;3xBRP averaged from measurements over 2-4 days, including sleep curves plotted in 30-min bins, daily sleep amount, sleep episode number, sleep episode duration and sleep latency at ZT12. Data from male flies are shown. n= 59-62. \**p* < 0.05; \*\**p* < 0.01; \*\*\**p* < 0.001; ns, not significant. Error bars: mean ± SEM.

## MATERIALS AND METHODS

### *Drosophila* stocks and maintenance

*w^1118^ iso31* was used as wild-type (*wt*), 2xBRP and background control. *brp* gene copy titration was achieved as previously reported (*12, 13*). Briefly, a *brp* null mutation (c04298, BDSC#85966) was utilized to reduce *brp* gene copy. A genomic *brp* P[acman] construct was used to increase brp gene copy number from 2xBRP to 3xBRP and 4xBRP. The *quiver/sleepless* alleles *sss^P1^* (BDSC#16588) and *sss^P2^* have been described previously (*8, 31*). *elav^C155^-Gal4* was used to express RFP-labelled Shaker pan-neuronally. Ectopic salivary gland expression was driven by *OK6-Gal4*. The following UAS lines were used in this study: *UAS-GFP-BRP* (*10*), *UAS-BRP* (*10*), and *UAS-Shaw-FLAG* (BDSC#55710) (*32*). The *UAS-Sh-RFP* line was generated by total synthesis and details can be found in the following sections.

Flies were reared under standard laboratory conditions and raised on semi-defined medium (Bloomington recipe) under a 12/12 h light/dark cycle with 65% humidity at 25°C. 5-to 8-day-old flies were used for all experiments. All fly strains were backcrossed to *w^1118^* (*iso31*, BDSC#5905) background for at least six generations or retained in an already isogenized background.

### Oxidative stress challenge assay

Survival under oxidative stress was performed as previously described (*13, 33*). Except for experiments related to the X-linked *Shaker mns* mutants in which only male flies were tested, female and male flies were separated and sorted into a population of 45-50 flies per vial at 2 days post-eclosion after fully mating. After 2 days, they were transferred into a vial containing 4 layers of filter paper soaked with 1.4 ml 5 % sucrose solution with 2 % H_2_O_2_. Flies were flipped into new vials with freshly prepared filter papers and solution every other day. The numbers of death were counted and recorded daily at the same time of the day. As results from female and male flies were essentially comparable, data from both sexes were pooled and comparison of survival curves was performed with GraphPad Prism using Log-rank analysis.

### Longevity

Longevity experiments were carried out exactly as previously reported (*13, 33*). Briefly, male flies were sorted into a population of approximately 20-25 flies per vial/replicate at the age of 2 days. To reduce the variability in life span, a few different cohorts of flies were used for each experiment. The longevity flies were regularly transferred onto fresh food every second day or over the weekend and the number of dead flies in each vial was recorded at each time of transfer until the death of the last fly of a replicate. Comparison of longevity curves was performed with GraphPad Prism using Log-rank analysis.

### Aversive olfactory learning and memory

Pavlovian aversive olfactory conditioning was performed as previous reported (*13, 33*). Briefly, two aversive odors, 3-Octanol (OCT) and 4-methylcyclohexanol (MCH), served as behavioural cues (odors were diluted in mineral oil at a 1:100 ratio), and 120 V AC current electrical shocks were used as a behavioural reinforcer. Briefly, during one training session, around 100 naive flies were placed in a T-maze and exposed to the first odor (conditioned stimulus, CS+, MCH or OCT) paired with electric shocks (unconditioned stimulus, US) for 60 s. Afterwards, these flies were allowed to rest for 60 s followed by exposure to the second odor (CS-, OCT or MCH) without electric shocks for 60 s. During testing session, these flies were simultaneously exposed to CS+ and CS-at the two sides of the T-maze and they had 60 s to choose between the two odors. A reciprocal experiment, in which the second odor was paired with electric shocks as CS+, was performed. A performance index was calculated as the number of flies choosing CS-minus the number of flies choosing CS+, divided by the total number of flies, and multiplied by 100 to get a percentage. The final performance index was the mean of the two performance indices from the two reciprocal experiments. STM was tested immediately after training, while 3h MTM was tested 3 h after training. As a component of 3h MTM, the 3h ARM was separated from 3h ASM by cold anesthesia. At 2 h 30 min after training, the trained flies received a cold anesthesia on ice for 90 s. 3h ARM was tested 30 min after the cold anesthesia, and 3h ASM was calculated by subtracting 3h ARM from 3h MTM.

### Sleep measurement

Sleep experiments were performed exactly as previously reported (*12, 13*). Briefly, sleep of single male flies was measured by Trikinetics Drosophila Activity Monitors (DAM2) from Trikinetics Inc. (Waltham, MA) in 12/12 h light/dark cycle with 65% humidity at 25°C. 3∼4-day-old single male flies were loaded into Trikinetics glass tubes (5 mm inner diameter and 65 mm length) which have 5% sucrose and 2% agar in one side of the tube. Flies were entrained for at least 24 h and their locomotor activity was measured at 1 min interval. Due to entrainment to new environment, data from the first ∼24 h were excluded. A period of immobility without locomotor activity counts lasting for at least 5 min was determined as sleep (*34*). Sleep from multiple days of recordings was analyzed and averaged using the Sleep and Circadian Analysis MATLAB Program (SCAMP) (*35*).

### Western blotting

6 adult brains per genotype were dissected in ice-cold haemolymph-like saline solution (HL3; 70 mM NaCl, 5 mM KCl, 10 mM NaHCO_3_, 20 mM MgCl_2_, 1.5 mM CaCl_2_, 5 mM HEPES, 5 mM Trehalose, 115 mM Sucrose, pH = 7.2) and stored in low-binding Eppendorf tubes on ice. Dissected brains were immediately centrifuged for 5 mins at 7000 rpm at 4°C, the HL3 supernatant was discarded, and the sample was frozen at -20°C. After completion of sample collection, 40 µl of lysis buffer (1x PBS, 0.5% Triton-X, 2% SDS, 1x Sample Buffer, 1x Roche cOmplete™ Mini protease inhibitor cocktail (#11836153001)) was added onto the brain pellets, and samples were homogenized by mechanical rotation for 1 h at 4°C, followed by 5 mins centrifugation at 13.000 rpm at room temperature. All genotypes/groups of an experiment were dissected on the same day, and sample collection for Western blotting was always performed in the middle of the day (around ZT5, i.e., 2 pm), to avoid confounding fluctuations in abundance of proteins under circadian control. Lysed brain samples were boiled for 5 mins at 95°C and subsequently centrifuged for 1 min at 13.000 rpm. An amount equal to 2 brains for each experimental condition was loaded into SDS-PAGE gels and immunoblotted according to standard protocols. Blots were manually developed with ECL solution and Kodak/GE films, and subsequently scanned by an EPSON V330 scanner and saved in 16-bit grayscale tiff format. The following primary antibodies were used: Rabbit anti-BRP^D2^ (rb5228, 1:40.000), rabbit anti-Shaker (Biozol PAB12776, 1:1000), mouse anti-Tubulin (Sigma T5168, 1:100.000). Secondary horseradish peroxidase (HRP)-conjugated goat anti-mouse and goat anti-rabbit antibodies were used for detecting the protein signals by chemiluminescence (Biozol, 1:5000).

### Sample preparation for label-free global proteomics

Sample collection and preparation for proteomic analysis was performed similar as for western blotting. 10 brains per genotype were hand-dissected in ice-cold HL3, centrifuged at 4°C and 7000 rpm for 5 mins, and immediately frozen at -80°C until completion of sample collection. Dissection was always performed in the middle of the day (around ZT5, i.e., 2 pm), and the 4 biological replicates were dissected on different days from independently collected fly cohorts from the same parental cross. 20 µl of lysis buffer was added onto each frozen brain pellet, and samples were lysed for 1 hour at 4°C on a rotator, followed by 5 mins centrifugation at 13.000 rpm. Subsequently, 1 μl of 100 mM Dithiothreitol (DTT) dissolved in 50 mM Triethylammonium-Bicarbonat (TEAB) buffer was added to the samples to reach a final concentration of 5 mM of DTT in the solution, and samples were incubated while shaking at 300 rpm for 30 mins at 55°C. Protein alkylation was initiated by addition of 2.1 μl of 400 mM Chloracetematide (CAA) dissolved in 50 mM TEAB, to reach a final CAA concentration of 40 mM inside the sample. Samples were then incubated while shaking at 300 rpm for 30 mins at room temperature in darkness. After alkylation, proteomic samples were boiled for 5 mins at 95°C, followed by 1 min of centrifugation at 13.000 rpm. Exactly 20 μl of each sample was loaded onto 4-20% Mini-PROTEAN® TGX™ precast gels (Bio-Rad laboratories, Hercules, USA), and SDS-PAGE was performed for 2-3 mins at 200 V until all proteins migrated out of the loading pockets into the gel. Coomassie staining of the gel was performed according to standard protocols, and each lane of protein sample was cut with sterile scalpels into 4 equal pieces. Gel pieces were transferred into low-binding Eppendorf tubes containing 50 µl of distilled H_2_O, and stored at 4°C until further processing.

### Liquid chromatography mass spectrometry (MS) analysis

In-gel protein digestion was carried out using trypsin at an enzyme-to-protein ratio of 1:20 (w/w) at 37°C overnight and 1 μg of peptide were loaded for the LC/MS analysis. The LC/MS analysis was performed using a Thermo Scientific Dionex UltiMate 3000 system connected to a PepMap C-18 trap-column (0.075 × 50 mm, 3 μm particle size, 100 Å pore size; Thermo Fisher Scientific) and an in-house-packed C18 column (column material: Poroshell 120 EC-C18, 2.7 μm; Agilent Technologies). After loading, peptides were separated at 250 nL/min flow rate over a 120 min gradient of increasing acetonitrile concentration and sprayed into an Orbitrap Fusion Lumos (instrument control software 3.1). The MS1 scans were performed in the Orbitrap. The MS2 scans were acquired in the ion trap with standard AGC target settings, an intensity threshold of 1e4 and maximum injection time of 35 ms. A 1 s cycle time was set between master scans.

### Data search and statistical analysis

Raw data was searched using MaxQuant (*27*) version 1.6.2.6 using standard settings. The number of missed cleavages allowed was set to 2, iBAQ and label-free quantification was enabled, and the match-between-runs option was disabled. The search was performed by target-decoy competition using the UniprotKB database of *Drosophila melanogaster* downloaded in May 2020 containing 42,678 entries. Results were reported at 1 % false discovery rate (FDR) for both peptide spectrum matches and the protein level.

Data analysis was performed using Perseus v 2.0.9.0 (*28*) and Python 3.10. Proteins identified only by site, reverse hits and potential contaminants were excluded. For further analysis, only protein detected in all 4 replicates of at least one genotype and in at least 8 out of 16 total replicates were kept. ANOVA with a permutation-based FDR correction of 5 % was used to compare the 4 groups, followed by post hoc Tukey’s HSD test. Hierarchical clustering based on Euclidean distance was applied to significant regulated hits. Data visualization and principal component analysis were performed using Python.

### Whole-mount *Drosophila* brain and salivary gland immunostaining

Brains of 3–5-day old female flies were used for the experiments. For salivary gland staining, tissues from third instar wandering larvae were collected. Adult files and larvae were dissected in ice-cold Ringer’s solution (130 mM NaCl, 5 mM KCl, 2 mM MgCl_2_, 2 mM CaCl_2_, 5 mM HEPES, 36 mM Sucrose, pH = 7.3). Dissected brains or salivary gland were fixed immediately in 4 % Paraformaldehyde (w/v) in PBS for 30–35 min on an orbital shaker at room temperature. Samples were washed in 0.7 % Triton-X (v/v) in PBS (0.7 % PBST) for 20 min × 3 times, followed by blocking in 10 % normal goat serum (v/v) in 0.7 % PBST for at least 2h at room temperature while shaking. After overnight (2 nights for mouse anti-BRP Nc82 and guinea pig anti-Shaker antibodies) primary antibody incubation at 4 °C, brains were washed in 0.7 % PBST for 30 min × 6 times at room temperature, followed by overnight secondary antibody incubation at 4 °C in darkness. Primary and secondary antibodies were diluted in 0.7 % PBST with 5 % normal goat serum. The following dilutions were used for the primary antibodies: 1:50 for mouse anti-BRP Nc82 (*10*), 1:500 for rabbit anti-GFP (*33*), 1:2000 for guinea pig anti-Shaker (*9*), 1:200 for rat anti-RFP (Chromotek, 5F8), 1:500 for guinea pig anti-BRP Last200 (*36*) and 1:200 for mouse anti-FLAG (*33*). A dilution of 1:300 was used for all secondary antibodies. After secondary antibody incubation and before mounting, samples were washed intensively with 0.7 % PBST for at least 30 min × 6 times. Then they were mounted on microscope slides in Vectashield and kept in a dark place at 4 °C before being scanned.

### Confocal microscopy and image analysis

Whole-mount adult brain samples were imaged on a Leica TCS SP8 confocal microscope (Leica Microsystems), and images were obtained using the Leica LCS AF software (Leica Microsystems). The central brain areas and salivary glands were imaged using a 40× and/or 63× oil objectives. For the same set of experiments, all the parameters were kept constant. Image stacks were processed and analyzed with Fiji (https://fiji.sc/). The quantification of fluorescence intensity within the central brain area was performed automatically via a custom-written Fiji Macro, with steps briefly as follows. Selected stacks containing regions interested were converted to a two-dimensional image using average Z-projection. Using one of the BRP neuropile staining channels as a “mask”, the central brain was segmented by the “Watershed Irregular Features” command from the BioVoxxel toolbox after background subtraction, image smoothing, and thresholding process. The average gray pixel value of the protein in question was measured inside this region.

### Generating UAS-Sh-RFP line

For Gal4-driven ectopic expression of Shaker in the larval salivary glands, the Shaker isoform F was synthesized and an RFP tag was incorporated into the MiMIC site of MI10885 (BDSC#56260). Plasmid cloning and microinjection were performed by WellGenetics Inc. The landing site of PBac{y[+]-attP-3B}VK00001 2R (59D3) was used for microinjection.

### Electrophysiology

Larval preparations and electrophysiology recording at *Drosophila* larval neuromuscular junction synapses were carried out as previously described with slight modifications(*12, 37*). Both two-electrode voltage clamp (TEVC) and single electrode current clamp recordings were performed on muscle 6 in the abdominal segments 2 and 3 of third instar larvae at room temperature. For dissection, larvae were pinned at their head and tail with their dorsal side facing up in modified Ca^2+^-free HL3 (70 mM NaCl, 5 mM KCl, 10 mM NaHCO_3_, 20 mM MgCl_2_ for TEVC/4 mM MgCl_2_ for current clamp, 5 mM HEPES, 5 mM Trehalose, 115 mM Sucrose, pH = 7.2), and then cut open dorsally along the rostro-caudal midline. The internal organs except for the central nervous system were removed without damaging body-wall muscles and pinning the larvae open with 4 additional pins. Recordings were then performed in 2 ml of HL3 used for dissection with CaCl_2_ added as indicated concentration in each experiment. For recordings containing 4-aminopyridine (4-AP), bath solution additionally contained 50 or 100 μM 4-AP or an equivalent volume of Milli-Q water and preparations were incubated for 5 minutes after being transferred into bath solution before recording. Preparations were visualized under a BX51WI Olympus microscope with a 40× LUMPlanFL/IR water immersion objective (Olympus Corporation, Shinjuku, Tokyo, Japan). Using sharp electrodes filled with 3M KCl, recordings were made from muscle cells with input resistances of ≥ 4 MΩ and initial membrane potentials between -30 and -80 mV depending on the external Ca^2+^ concentration. Two cells were recorded per animal. The recording electrodes (20-50 MΩ) were pulled using a Flaming Brown Model P-97 micropipette puller (Sutter Instrument, CA, USA). The stimulation electrodes were pulled using a DMZ-Universal-Electrode puller (Zeitz-Instruments Vertriebs GmbH, Martinsried, Germany), then tip openings were shaped by fire-polishing with a DMF1000 microforge (World Precision Instruments Inc., FL, USA). Recordings were acquired using an Axoclamp 2 B amplifier with HS-2A x0.1 head stage (Molecular Devices, CA, USA) and sampled at 10 kHz with an Axon Digidata 1322 A digitizer (Molecular Devices, CA, USA). Signals were low pass filtered at 1 kHz using an LPBF-48DG output filter (NPI Electronic, Tamm, Germany).

Miniature junction currents (mEJCs) were recorded for 90 seconds with the voltage clamped at -80 mV; for all other TEVC recordings were performed while clamping the voltage at -60 mV. Single evoked excitatory junction currents/potentials (eEJCs/eEJPs) were recorded after stimulating the cut end of the segmental motor neuron bundle with 5 V, 300 μs at 0.2 Hz using an S48 Stimulator (Grass Instruments, Astro-Med, Inc., RI, USA). Paired pulse recordings were performed using the same stimulation twice at 10 or 30 ms interval. For 10 Hz trains, a single train was recorded for 1 s with 100 stimulations in total. To estimate the size of readily releasable synaptic vesicle pool (RRP), single train recordings with 61 stimulations were performed at 100 Hz.

The recordings were analyzed with pClamp 10 (Molecular Devices, Sunnyvale, CA, USA), GraphPad Prism 6 (GraphPad Software, Inc) and two custom-written Python scripts utilizing the pyABF package for Python 3.10 (*38*). Briefly, mEJCs were filtered with a 500 Hz Gaussian low-pass filter, then analyzed using the template search function of the pClamp 10 software. The eEJCs/eEJPs traces were analyzed using custom-written Python scripts, as described below. An average trace was generated from 20 eEJCs/eEJPs traces per cell for single 0.2 Hz stimulation and from 10 traces for 10 ms/30 ms interval paired pulse recordings. Stimulation artifacts of eEJCs/eEJPs were removed for clarity. Decay constant τ was calculated by fitting a first order decay function to the region of the average trace of the single 0.2 Hz stimulation recording from 60% to 5% of the total amplitude after the peak. Charge was calculated by integrating the traces along the baseline using the composite trapezoidal rule. Paired pulse ratios were calculated by dividing the amplitude after the second pulse by the amplitude after the first pulse. Estimation of the RRP size and refilling rates were performed as described previously (*39*). The amplitudes of the eEJCs in the 100 Hz train were extracted from the recording. For each cell, these amplitudes were then added cumulatively and divided by the average mEJC amplitudes as measured for this cell to obtain the cumulative quantal content. This cumulative quantal content was then plotted against the number of stimulations and a linear regression was performed for the last 20 amplitudes, where the cells reach a steady state due to the depletion of the RRP. The intersect of this linear fit with the Y-axis is the estimated RRP size, the slope of the linear fit is the estimated refilling rate.

For mean-variance analysis, the TEVC recordings were performed in the bath solution initially contained 0.375 mM Ca^2+^ and after each measurement, 1 ml of bath solution was carefully removed with a pipette while the cell was still clamped. Then, 1 ml of HL3 containing an appropriate concentration of Ca^2+^ to adjust the Ca^2+^ concentration of the bath solution to twice its original value (0.375 → 0.75 → 1.5 → 3 → 6 mM Ca^2+^) was carefully added. The bath solution was then very gently mixed, and the cell was allowed to adjust to the new external Ca^2+^ concentration for 1 minute before the next recording. Quantal size *q*, release site number *N* and release probability *p* were determined from recordings obtained from the same cell at different Ca^2+^ concentrations based on the theoretical considerations of Clements and Silver (*40, 41*). Briefly, at each Ca^2+^ concentration, the average amplitude of eEJCs traces for each cell was plotted against the variance of their amplitudes to obtain a parabola fit. The standard equation for a parabola is *y* = *Ax* − *Bx*^2^ (1), where *y* is the variance of the amplitude, *x* is the mean amplitude, and *A* and *B* are the fitting parameters of the amplitude. Assuming that the coefficient of variation of the eEJCs amplitudes at individual release sites is negligible, quantal size *q* was estimated as *q* = *A* (2). A lower limit for the number of release sites *N* was then calculated from this parabola fit as 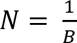. Using the same assumption as in (2), release *p* probability *p* was then individually calculated for each Ca^2+^ concentration as 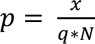.

## Acknowledgments

We thank Kyunghee Koh, Bloomington *Drosophila* Stock Center (BDSC), Harvard Medical School, and Vienna *Drosophila* Resource Center for fly lines. We are grateful to Kyunghee Koh and Developmental Studies Hybridoma Bank for antibodies. This work was supported by grants from European Research Council (ERC) advanced grant (SynProtect 101097053) and the Deutsche Forschungsgemeinschaft (DFG; CRC1315/A08 Project ID 327654276) to S.J.S. S.H. was supported by the Leibniz SAW SyMetAge and Charité NeuroCure. NeuroCure is funded by DFG under Germany’s Excellence Strategy -EXC-2049 -390688087. C.P. was supported by DFG under FOR5228/RP8 (Project ID 447288260) and NeuroNex (Project ID 436260754).

## Author contributions

Conceptualization, data interpretation and writing: S.J.S.

Investigation: S.H., C.P., M.E., D.T., C.B.B., T.G., N.R., Z.Z., O.T., A.M.W., and F.L.;

Funding acquisition: S.J.S.

